# Cooperative breeding in a plural breeder: the vulturine guineafowl (*Acryllium vulturinum*)

**DOI:** 10.1101/2022.11.23.517633

**Authors:** Brendah Nyaguthii, Tobit Dehnen, James A. Klarevas-Irby, Danai Papageorgiou, Joseph Kosgey, Damien R. Farine

**Affiliations:** Department of Ornithology, National Museums of Kenya, P.O. Box 40658-001000, Nairobi, Kenya; Mpala Research Centre, P.O Box 555-10400, Nanyuki, 10400, Kenya; Department of Collective Behavior, Max Planck Institute of Animal Behavior, 78464 Konstanz, Germany; Centre for Ecology and Conservation, University of Exeter, Penryn Campus, Penryn TR10 9FE, UK; Department of Evolutionary Biology and Environmental Studies, University of Zurich, 8057 Zürich, Switzerland; Department of Migration, Max Planck Institute of Animal Behavior, Radolfzell, Germany; Department of Biology, University of Konstanz, Konstanz, Germany; Kenya Wildlife Service, P.O. Box 40241-001000, Nairobi, Kenya; University of Eldoret, School of Natural Resource Management, Department of Wildlife, 1125-30100 Eldoret, Kenya; Division of Ecology and Evolution, Research School of Biology, Australian National University, 46 Sullivans Creek Road, Canberra, ACT 2600, Australia

**Keywords:** Cooperation, Galliformes, helping, precocial breeding, vulturine guineafowl

## Abstract

Cooperative breeding is widely reported across the animal kingdom. In birds, it is hypothesised to be most common in altricial species (where chicks are dependent on parental care in the nest after hatching), with few described cases in precocial species (where chicks are more independent immediately after hatching). However, cooperative breeding may also be more difficult to detect in precocial species and therefore has been overlooked. In this study, we investigate whether vulturine guineafowl (*Acryllium vulturinum*)—which have precocial young—breed cooperatively and, if so, how care is distributed among group members. Using data collected from colour-banded individuals in one social group of vulturine guineafowl over three different breeding seasons, we found that multiple females can attempt to reproduce in the same breeding season. Broods had close adult associates, and most of these associates exhibited four distinct cooperative breeding behaviours: babysitting, within-group chick guarding, covering the chicks under the wings and calling the chicks to food. Further, we found that offspring care is significantly male-biased, that non-mother individuals provided most of the care each brood received, that breeding females differed in how much help they received, and that carers pay a foraging cost when providing care. Our results confirm that vulturine guineafowl are cooperative breeders, which they combine with an unusual plural-breeding social system. Our study also adds to growing evidence that cooperative breeding may be more widespread among species with precocial young than previously thought, thereby providing a counterpoint to the altriciality-cooperative breeding hypothesis.

## INTRODUCTION

In some species of birds, mammals, fish and invertebrates more than two adults contribute towards raising offspring, which is known as cooperative breeding. Typically, groups of cooperatively breeding individuals consist of a breeding pair and their offspring from previous breeding attempts that delayed dispersal and provide offspring care to their younger siblings (Clutton-Brock, 2002). As cooperative breeding seems to be a suboptimal reproductive strategy for such individuals, it has received considerable empirical (Koenig & Dickinson, 2016; Stacey & Koenig, 1990) and theoretical (Emlen, 1982; Hatchwell & Komdeur, 2000; Shen et al., 2017) attention over the past five decades. In birds, cooperative breeding is thought to be more prominent among altricial species (Cockburn, 2006; B. Wang et al., 2017). A recent review found that while 11% of species with altricial young breed cooperatively, cooperative breeding is found in only 4% of species with precocial young (Scheiber et al., 2017). The relative paucity of cases among species with precocial young has led to the hypothesis that there exists a link between altriciality and cooperative breeding, because precocial chicks require less care (Ligon & Burt, 2004; N. Wang & Kimball, 2016). Yet, early studies on species with precocial chicks, such as the pukeko (*Porphyrio melanotus*), played an important role shaping the field of cooperative breeding (Craig & Jamieson, 1990). Further, several new species with precocial young have recently been added to the list of cooperative breeders, including the Kalij pheasants (*Lophura leucomelanos*) (Zeng et al., 2016) and the buff-throated partridge (*Arborophila brunneopectus*) (B. Wang et al., 2017). Thus, cooperative breeding has likely been generally under-explored among precocial breeders.

The disparity in the presence of cooperative breeding between altricial and precocial species could be because species with precocial offspring—which are more independent straight after hatching—have less need for care. However, this logic may be flawed. Due to the high predation risk to the eggs of ground-nesting birds (Thompson & Raveling, 1987), females of ground-nesting precocial species typically have very high nest attendance rates, thus foregoing feeding during incubation. For example, female ring-necked pheasants (*Phasianus colchicus*) increasingly attend their nest throughout laying and, as a result, suffer a reduction in body mass equating to approximately 19% (Breitenbach & Meyer, 1959). Given the substantial costs associated with egg laying and incubation to breeding females, reduced post-hatching costs could allow breeding females to recover from their reproductive investment and, thereby, produce more chicks (the load lightening hypothesis) (Crick, 2008; Hatchwell, 1999). Furthermore, given that chicks of most precocial species have lower survival probabilities than adults, additional offspring care could also enhance chick survival in such species (Heinsohn, 2004), albeit likely to a lesser degree than in altricial species. Accordingly, there is substantial scope for non-breeding individuals to gain sufficient indirect fitness by reducing the effort of related females.

An alternative reason for the limited evidence for cooperative breeding in precocial species is that classical examples of cooperative breeding in birds primarily consider offspring care at the nest, rather than other forms of care that may occur after chicks leave the nest. If this is the case, then cooperative breeding may have been disproportionately undetected in precocial species. While provisioning chicks at the nest provides parents and/or chicks with obvious benefits, there are many other ways in which individuals can contribute to raising offspring that are also possible in precocial species. For example, in trumpeters (*Psophia spp*.), which have precocial offspring, non-breeders contribute to nest building and incubation of eggs, as well as providing chicks with food after leaving the nest (Sherman, 1995). Even without attending the nest, group members could still enhance offspring survival, or reduce parental investment, post-hatching. For example, group members may protect chicks from predators or the abiotic environment (such as providing chicks with shade or warmth), identify and provide food for chicks, or maintain vigilance to allow chicks more time to forage. Such forms of cooperative breeding, however, are likely more difficult to detect and thus require more careful observations than those necessary to describe offspring care at the nest. Beyond the challenge of quantifying the benefits to chicks and costs to individuals providing care, it can also be difficult to study precocial chicks in their natural environment, as this requires following groups and chicks closely for long periods, during which they may be inaccessible to researchers. Additionally, social groups generally need to be habituated to allow close observations—thereby increasing the investment required to study cooperative breeding in species with precocial young. Species in which offspring care is primarily provided away from the nest may thus be less likely to be studied for their cooperative breeding behaviour. Together, these factors could have led to a general under-reporting of cooperative breeding in species with precocial young.

One group of birds with precocial young in which evidence for cooperative breeding is increasing is Galliformes. To date, support for the existence of cooperative breeding has been found in nine species of Galliformes. In five species, the buff-throated partridge (*Arborophila brunneopectus*) (B. Wang et al., 2017), black-breasted wood quail (*Odontophorus guttatus*) (Hale, 2006), Kalij pheasants (*Lophura leucomelanos*) (Zeng et al., 2016), Tibetan eared pheasants (*Crossoptilon crossoptilon*) and white-eared pheasants (*Crossoptilon harmani*) (Lu & Zheng, 2005), the evidence for cooperative breeding is relatively strong. In these species, group members were found to show food to the chicks, remain vigilant against predators and care for the chicks by expressing agonistic interactions among conspecific intruders. In four species, the California quail (*Callipepla californica*) (Lott, 1999), northern bobwhite (*Colinus virginianus*) and scaled quails (*Callipepla squamata*) (Orange et al., 2016) and marbled quail (*Odontophorous gujanensis*) (Skutch, 1947), the evidence remains largely anecdotal. The phylogenetic distance between these species, and among other species that breed cooperatively and have precocial young, suggests that cooperative breeding could be more common in Galliformes—and in precocial species more generally—than previously thought. This is likely because Galliformes have received little attention from researchers relative to many other groups of birds, especially in the context of cooperative breeding. Thus, Galliformes represent an excellent taxonomic group in which to explore cooperative breeding, and how cooperative breeding behaviours might be expressed away from the nest.

The vulturine guineafowl is a large, terrestrial Galliforme that lives in stable groups of approximately 13-65 individuals, with groups associating preferentially with other groups to form a multilevel society (Papageorgiou et al., 2019). Given that social groups contain many adults, sub-adults and juveniles (Papageorgiou et al., 2019), vulturine guineafowl are a good candidate for being both plural and cooperative breeders, the former being defined as living in stable social groups containing multiple reproductive males and females. Vulturine guineafowl also have extreme sex-biased dispersal—all males remain in their natal groups (Klarevas-Irby et al., 2021)—setting the scene for high within-group relatedness among males, which could drive indirect fitness benefits of cooperative breeding (Hamilton, 1964). Furthermore, a recent comparative analysis suggests that the same environmental conditions might promote both cooperative breeding and multilevel societies (Camerlenghi et al., 2022). The main focus of our study is to characterise the incidence and structure of cooperative breeding in the vulturine guineafowl. We quantify the association between each member of the group and each brood. We also test what share of offspring care is given by mothers versus other group members, and whether the latter pay a foraging cost when providing care. By employing a heterogenous approach to our study of cooperative breeding in the vulturine guineafowl, we are able to provide new evidence for the presence of cooperative breeding behaviour in an under-represented group of precocial species.

## METHODS

### Study species

Although they live in large groups with stable membership for most of the year, vulturine guineafowl form pairs at the beginning of the breeding season (Papageorgiou et al., 2021), after which pairs move separately from the remaining group and the male mate-guarding the female. This is followed by the female laying—and independently incubating—a clutch of 13-15 eggs in a scrape on the ground (Del Hoyo et al., 1994). (We note that our field observations indicate that clutch sizes are smaller, and more variable, than the literature suggests, ranging from 7 to 12 eggs across 14 nests found). While the female incubates, the male re-joins the group, and can re-pair with a new female. Vulturine guineafowl chicks are precocial, with the mother and chicks typically re-joining the social group (or part of their social group) very soon after hatching. As in other guineafowl species (Del Hoyo et al., 1994), vulturine guineafowl females appear to receive no help during incubation. Although information about this remains anecdotal, we have never observed evidence suggesting otherwise (e.g., via GPS tracking and a few nests monitored using camera traps). Chicks are highly vulnerable to predation during the first few weeks of life, meaning that they could benefit from protection offered by other group members.

### Study group and field data collection

Our study is focused on a long-term habituated social group living mostly within the fenced area of the Mpala Research Centre (MRC), in Laikipia county, Kenya, and forms part of a long-term study population of vulturine guineafowl at the MRC. Mpala is characterised by semi-arid savanna habitat with rainfall averaging between 500 and 600 mm per year, occurring predominantly in two rainy seasons (Young et al., 2003). The natural vegetation is mainly *Acacia* scrubland, and vulturine guineafowl specialise on the red soils that are dominated by *Acacia mellifera* and *Acacia etbaica*. The study group was first colour-banded, and tracked with GPS (He et al., 2022), in September 2016. Group size over the study period ranged from 30-35 adults. In the wet seasons spanning April to May 2019, April to May 2020, and October to November 2020, we recorded breeding activity within the study group. This involved following pairs that split from the main group, finding and monitoring the nests, and observing care behaviours post-hatching. Pairs were defined as a female and an associated male who moved together (that is, typically less than 5 m apart) and away from the group (that is, more than 20 m from other group members, but generally much further). Before considering two birds to be a pair, they had to be observed moving together for the whole day, but all pairs were recorded even if they were paired only for a single day. By following pairs, we eventually located and monitored the nests of all breeding females. When the clutch was expected to hatch (25 days after the start of incubation), we searched for the female and chicks.

We also collected data on associations between group members and broods by recording group composition data in the morning (0600–0930 hrs) and evening (1700–1900 hrs) for an average of four days per week. Each time members of the group were encountered, the identity of every adult bird present was recorded, as well as the total number of banded and unbanded adults, and the number of chicks present from each of the broods. All birds that were present together in an area (that is, within sight of each other and behaving cohesively) were considered to be part of the same group, and given the same group identifier. Within these individuals, if some were more spatially clustered (e.g., within two metres of each other but more than 10m from other individuals), then these were recorded as a subgroup within the broader group (or as pairs, if they remained distinctly separate for the entire day).

Although chicks could not initially be marked individually, broods (up to N=3 per season) were generally identifiable because they hatched asynchronously, meaning that there was a clear size difference between chicks from different broods. From these differences, we ascertained that those chicks from the same brood remained together the vast majority of the time. However, there were two exceptions. During two seasons, multiple broods merged after hatching at about the same period. We therefore found a corresponding increase in overlap among the adults associated with each brood over the second and third season, but we cannot know whether the patterns we observed are common or not when broods are very close in age. Association data, between adults and broods, were collected for each breeding season until chicks were four months old.

In addition to collecting association data, we observed and recorded cooperative breeding interactions via all-occurrence sampling (Altmann, 1974) over the three seasons. Specifically, we found four types of cooperative breeding interactions to occur between adults and chicks. The first is babysitting behaviour (BBS), whereby individual A stays more than 20 metres from the rest of the group members with one or more chicks. The second is within-group chick guarding behaviour (GRD), whereby an individual does not allow other adult group members to approach one or more chicks. The third behaviour is chick covering (COV) (Figure 1A), whereby an individual covers one or more chicks under its wings. The final cooperative breeding behaviour is chick feeding (CFD) (Figure 1B), whereby an individual performs soft trills (a type of vocalisation) calling chicks to a food item. For each interaction, we recorded the identity of the actor, the identity of the recipient brood, the event duration in the case of cover events, and the number of chicks involved in each interaction. Because broods often moved separately through the group’s home range, we aimed to distribute our observation effort evenly across broods over each season.

**Figure 1.**
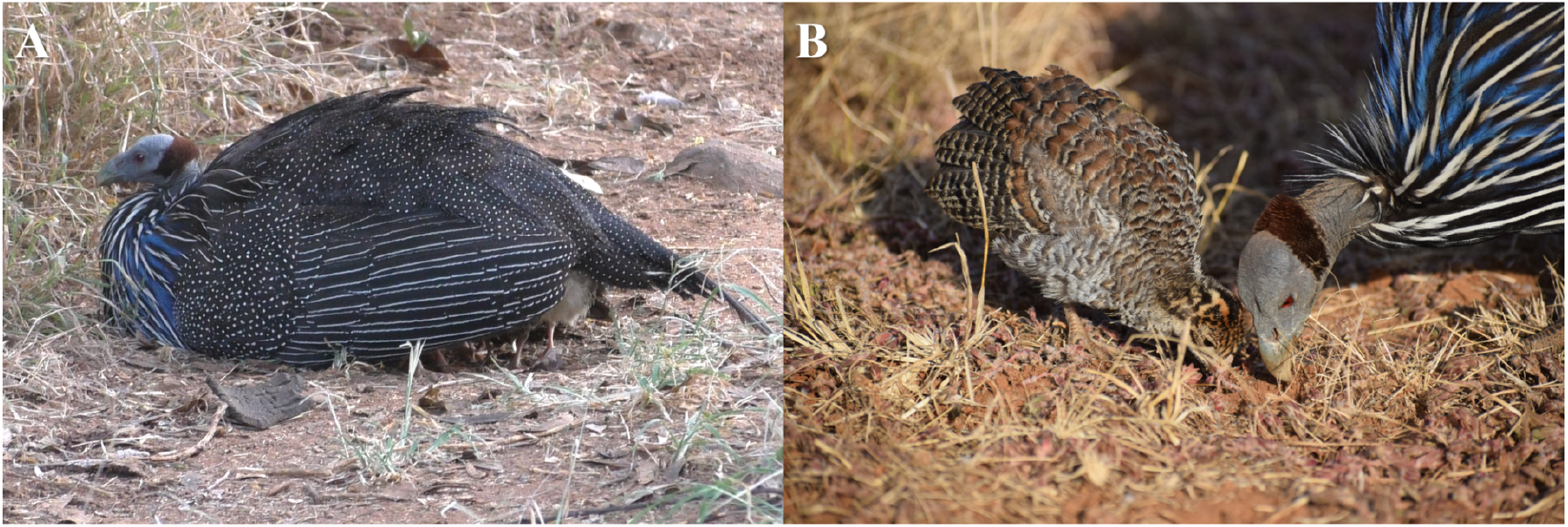
Examples of cooperative breeding behaviours. (A) Chick covering behaviour by an adult male and (B) chick feeding behaviour by an adult male.

We defined any individual that engages in one or more of the four cooperative breeding behaviours described above as a ‘carer’. These carers can include non-parents (e.g. older siblings from previous years (Ligon & Ligon, 1978), unrelated individuals (Clutton-Brock, 2002) or parents (both genetic and social). Due to a lack of data on paternity, we can only conclude which female is the social mother. However, from our monitoring of the study group, we know that many carers are the older (social) offspring of the female whose chicks they provided care to, and that some individuals in our study engaged in cooperative breeding behaviours prior to reproductive maturity (estimated age of maturity is 2 years (Del Hoyo et al., 1994), and males only reach the size of adults after more than 12 months). Thus, for each bird in each season, we provide three levels of evidence that point to some individuals being non-breeding helpers: (1) individuals that provide care prior to being sexually mature, (2) individuals that provide care to the brood of their social mother, and (3) individuals that provide care prior to ever having been observed forming a pair themselves.

Finally, to quantify whether carers pay a cost by engaging in cooperative breeding behaviour, we recorded videos during a period when the chicks were younger than five weeks (in the final season). From these videos, we were able to quantify the foraging activity by birds in each recording session for the duration that an individual could be tracked in the frame without moving out of the frame (due to occlusion or due to the movement of the person holding the camera). A new session would start if the focal individual started covering the chicks under the wings and would end when they stopped performing the behaviour. Each time we recorded an individual covering the chicks, we also recorded another focal individual who was not engaged in cooperative breeding behaviour. For each focal observation, we counted the number of pecks the individual made (pecking on the ground or on a plant) divided by the length of the focal observation, and whether or not the individual was covering the chicks under its wings or not.

### Data analysis

#### Who cares and who receives the most care?

We used a social network approach to identify the main carers for each group in each of the three seasons. Using the group composition data, we calculated the rate of attendance of each group member to each brood. This rate was calculated by dividing the number of subgroups that the individual was observed in that comprised the brood by the total number of subgroups that the individual was observed in, limited to group composition observations containing that brood (because not all broods were observed in a given sampling day). For example, if an individual was observed 10 times in different groups that contained brood A, of which 8 times it was in a subgroup with brood A, the rate of attendance of the individual to brood A was 0.8.

To identify which individuals occurred with the brood more than expected by chance we used a simple permutation test (Farine, 2017), which consisted of randomising the subgroups that the focal chicks were contained in. The permutation test worked as follows: for each day, K observations of the chicks were randomly allocated to subgroups observed on that day, where K corresponds to the number of subgroups the chicks were originally observed in that day (thus, all potential carers remained in the same subgroup, but the chicks were moved between subgroups for the purpose of the permutation test). The rate of attendance for each individual was re-calculated from these permuted data. The permutation test therefore maintained the number of times each individual and each brood was observed, and the same number of total groups and subgroups. This permutation procedure was repeated 1000 times, thereby generating a distribution of the rate of attendance values for each individual. From this distribution, we extracted the identities of the individuals whose observed rate of attendance was higher than 95% of the rates generated by the permutation procedure (i.e., significant at P≤0.05 using a one-tailed test). These individuals were recorded as having significantly higher attendance to that brood than expected by chance. This process was repeated for all broods in all seasons. Finally, to test whether males were disproportionately represented as carers, we conducted a two-sample proportion test that compared the proportion of males to females among the carer and non-carer categories (excluding mothers) in each of the three seasons.

We used the cooperative breeding interaction data to characterise the relative contribution of each significant associate to the brood. From these data, we determined whether the mother provided most of the care or not, and (if not) how help received varied between broods. First, we counted the number of each interaction directed towards a focal females’ chicks. Because babysitting and within-group guarding behaviours were expressed relatively rarely, they were combined into one category for analysis. Mothers clearly did not give the majority of the care to their chicks, so we then tested whether some females received more help than others in each of the three seasons. As we did not have an equal opportunity to observe each brood, and difference can lead to spurious outcomes (Hoppitt & Farine, 2018), we used a two-sample test for equality of proportions. We thus conducted pairwise contrasts of the proportion of total help each mother gave to the chicks in her brood. This comparison was performed across the broods within each of the three seasons. Significant effects mean that the difference in the proportion of help given by the mother was significantly lower in one brood than the other.

Finally, we tested whether males gave overall more help than females. To do this, we counted the number of chick-feeding interactions for each individual in each season as they were the most prevalent compared to other interactions. We then constructed a generalised linear mixed model with a Poisson error distribution, sex as the only predictor of the count of chick feeding interactions per individual per season, and individual identity and season as nested random effects. We excluded mothers from this analysis. While we could not explicitly account for differences in the number of observations, in practice every individual had the same opportunity to be observed helping each time we collected data. Thus, under random feeding, we would not expect our model to produce a significant difference.

#### Do non-mother individuals pay a cost for caring?

Using the data extracted from the videos, we quantified the proportion of the time spent foraging (response variable) and tested whether this was predicted by whether the focal individual was involved in the COV (wing-covering) behaviour (1) or not (0) (binary independent variable). To do this, we used a general linear model with a binomial error distribution. Ideally, we would have accounted for repeated observations of some individuals using mixed models, but we unfortunately could not always identify the individual recorded, and on only three occasions could we do so while an individual was covering the chicks. However, the data were relatively well distributed amongst individuals (average 2.25 observations per individual, when the identity was known). We thus do not expect repeated observations to have a major impact on our conclusions.

## RESULTS

### Multiple females breed in groups of vulturine guineafowl

Multiple female group members bred in each of the three breeding seasons (Table 1). Among these, there was some consistency in which females attempted to breed, with two females attempting to breed in all three seasons. Others were only observed breeding in later seasons, suggesting that they might have been recently recruited into the group and we observed their first reproduction. Overall, it appears that reproduction is open to all female group members, with the exception of females that have not yet dispersed.

**Table 1:** Females in each of the three seasons that successfully bred (had chicks) and the number of significant associates with the brood if they successfully hatched (Y=Yes, N=No).

| Season 1 (May 2019) |  |  | Season 2 (May 2020) |  |  | Season 3 (Nov 2020) |  |  |
| --- | --- | --- | --- | --- | --- | --- | --- | --- |
| Female | Successful | Number of associates | Female | Successful | Number of associates | Female | Successful | Number of associates |
| BYYO<sup>1</sup> | N | NA | GOOP | Y | 28 | GAGA | Y | 10 |
| GOBO | Y | 3 | MRMW<sup>4</sup> | N | NA | GOOP | Y | 10 |
| GOOP<sup>2</sup> | Y | NA | RARA<sup>5</sup> | N | NA | MGYB<sup>8</sup> | N | NA |
| RGBW | N | NA | RGBW | Y | 28 | MRMW<sup>9</sup> | N | NA |
| ROOR<sup>3</sup> | Y | NA | WOBY<sup>6</sup> | N | NA | RKGK | Y | 14 |
| WOBY | Y | 6 | WYPO<sup>7</sup> | N | NA | YOBK | Y | 7 |
| YOBK | Y | 7 | YOBK | Y | 11 |  |  |  |

<sup>1</sup>BYYO was depredated during incubation.
<sup>2</sup>GOOP's chicks were all depredated by the second week.
<sup>3</sup>ROOR was depredated the first night after re-joining the group. <sup>4</sup>MRMW's nest was depredated by a white-tailed mongoose. <sup>5</sup>RARA dumped her eggs on MRMW's nest, which was later depredated. <sup>6</sup>WOBY was depredated during incubation. <sup>7</sup>WYPO's nest was depredated during incubation. <sup>8</sup>MGYB's nest was depredated on the third day of the egg-laying period. <sup>9</sup>MRMW's nest was depredated after she abandoned it during incubation.

### Carers are predominantly male, and include non-breeders

There were four key insights from our analyses of significant associates (Figure 2). First, each brood had a number of individuals that were observed with the brood more than expected by chance, and not all individuals were consistently detected with a brood. Second, there was relatively little overlap amongst the individuals that were significantly associated with each brood, indicating that each brood had a distinct set of associates. Third, there was a significant male bias among associates (Table 2), and out of all the significant associates only one non-breeding female was observed providing care (BYBY in the first year, who was a subadult female). Finally, multiple lines of evidence suggests that a number of significant associates (and individuals that provide care) are non-breeding helpers, including individual males that (1) were not sexually mature at the time of helping (and not fully grown), (2) were helping their social mothers, and (3) helped prior to ever having been observed forming a pair with a female.

**Figure 2.**
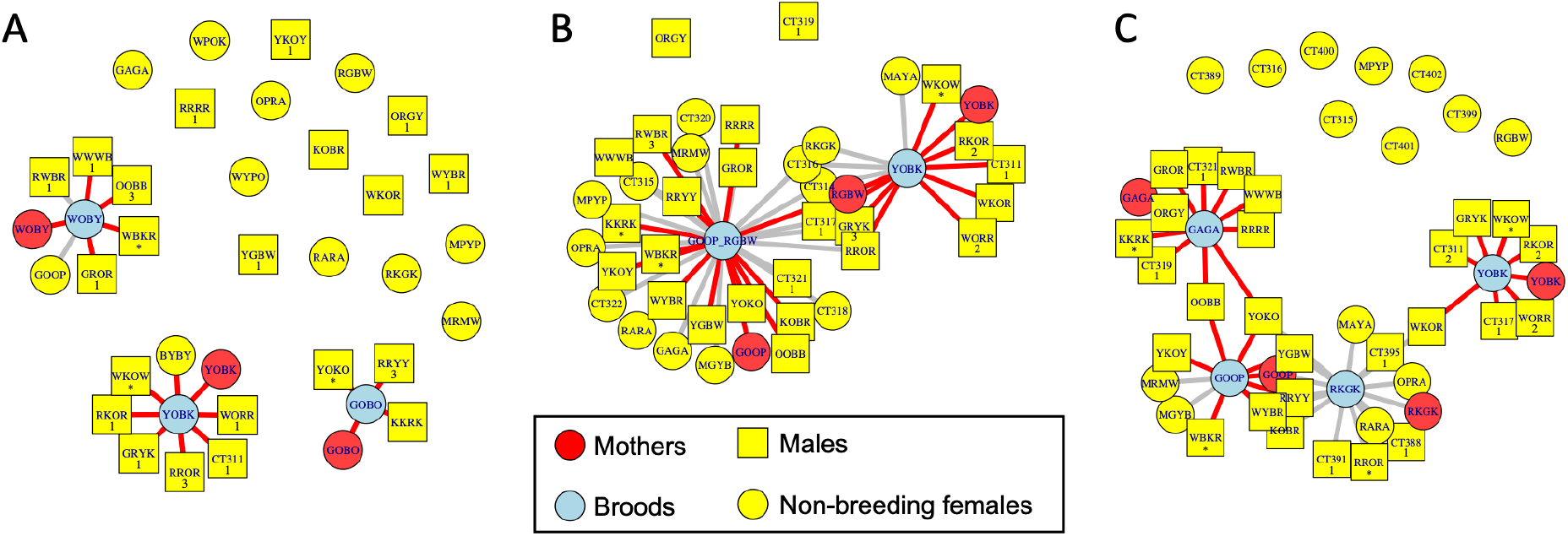
Social networks of attendance to the broods in (A) May 2019, (B) May 2020 and (C) Nov 2020 breeding seasons. Red edges represent significant associates who were also observed engaging in caring behaviours. Numbers represent levels of evidence for males being helpers as: (1) caring before being sexually mature, (2) caring for the chicks of their social mother, (3) caring before being observed engaging in any reproductive behaviours. For example, the male CT311 was only six months old in the first season (May 2019), and, at that time, was less than 50% the size of an adult male (∼800g vs 1.8kg). Males marked by a * were observed paired with the female of the broods that they associated with significantly. ROOR was depredated and her chicks joined the WOBY brood in May 2019, while the broods of GOOP and RGBW merged during the May 2020 season. Each of the non-significantly associated males in May 2019 were observed giving care at least once, but none of the non-significantly associated males in May 2020 were observed giving any care.

**Table 2.** Males are significantly more associated with the broods than females. Results of the two-sample test for equality of proportion of males among the non-associated individuals versus males among the significant associates of the three broods, in each of the three seasons.

| *Proportion of significant associates* |  |  |  |  |  |  |
| --- | --- | --- | --- | --- | --- | --- |
| *Season* | *Male* | *Female* | *X<sup>2</sup>* | *DF* | *P* | *95%CI* |
| 1 | 0.667 | 0.182 | 4.987 | 1 | 0.023 | 0.111 – 0.858 |
| 2 | 0.815 | 0.889 | 0.063 | 1 | 0.801 | -0.327 – 0.178 |
| 3 | 1 | 0.455 | 13.63 | 1 | 0.0002 | 0.187 – 0.904 |

### Males provide most care while mothers provide a small proportion of the total care to the offspring

Mothers also consistently provided a small proportion of the total care that their offspring received. For example, in the first season, out of the 20 cover events recorded for YOBK’s chicks, the mother was found to cover chicks in one event for only one minute. She also contributed only 42 of the 330 chick feeding events and one out of seven between group guarding and within-group guarding events. However, not all females received the same amount of help from carers. YOBK, a female who is known to have successfully bred before, generally received more help in food provisioning and covering chicks under the wings than the other females (see Supplementary Tables S1–S3), whereas there were fewer differences in the help received among the other mothers (see Supplementary Tables S4–S9). YOBK received more help with chick provisioning in each of the other two seasons, with the exception of one female (GAGA) in the third season (see Supplementary Table S7). As cover interactions were relatively rare, some comparisons could not be drawn effectively. There were no instances of babysitting and guarding behaviours in the first season for two mothers as well as the third season for all the mothers.

Overall, males provided significantly more caring interactions than females. Of the 2506 chick feeding interactions given by non-breeding individuals that we recorded across the three seasons, 2451 were given by males. This translates to an estimated 32.7 times more interactions per individual per season for males relative to females (Table 3).

**Table 3.** Males provide significantly more care than females. Results of the generalised linear mixed model comparing the number of chick-feeding interactions performed by male and female non-breeders across the three seasons.

|  | *Coefficient* | *SE* | *Z* | *P* |
| --- | --- | --- | --- | --- |
| Intercept | -0.703 | 0.569 | -1.235 | 0.217 |
| Sex (M) | 3.487 | 0.507 | 6.880 | <0.001 |
|  | *Random effect* | *Variance* | *SD* |  |
|  | Actor | 2.146 | 1.465 |  |
|  | Season | 0.433 | 0.658 |  |

### Non-mother individuals pay foraging costs while caring

Based on 27 observations of covering chicks under the wings, where cover duration data were available, individuals performed this behaviour on average for 16 minutes (range=1–60 minutes). From the video data, we estimated that birds pecked on average 0.10 times per second when they were not performing the cover behaviour (range=0–1.47) and pecked on average 0.01 times per second when they were performing the cover behaviour (range=0–0.01), a significant difference (Table 4). This means that, on average, a bird performing a key caring behaviour missed 81 pecks (90 pecks over 16 minutes when not helping versus 9 pecks over 16 minutes when helping), thereby potentially reducing their foraging intake (based on pecks) by 90%.

**Table 4.**
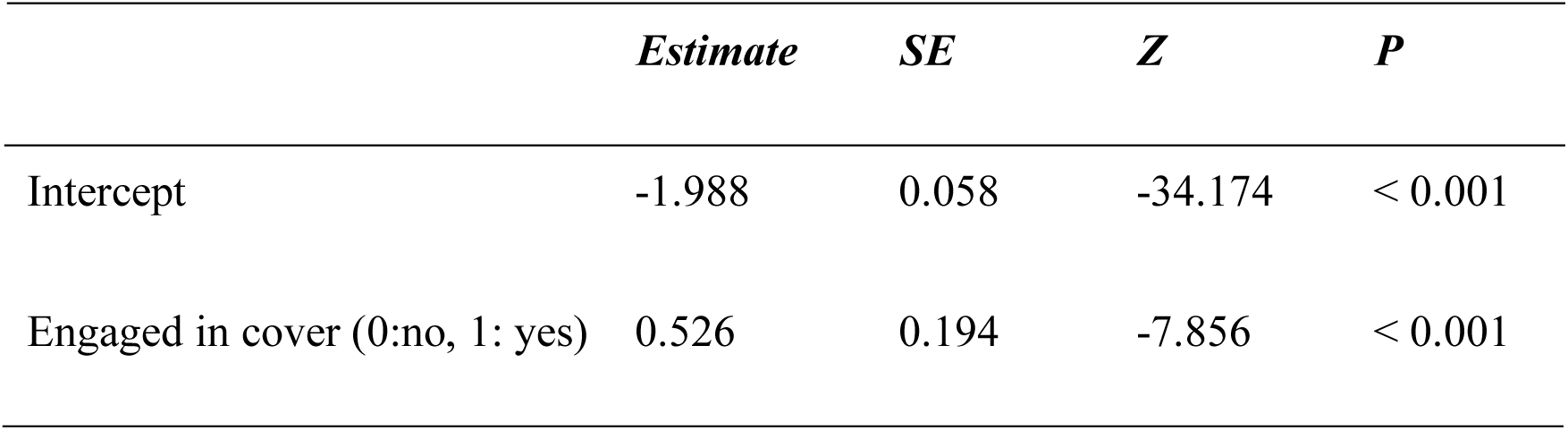
Non-mother individuals pay a foraging cost when caring. The proportion of the time spent foraging (response variable) was predicted by whether the focal individual was involved in chick covering behaviour. The estimates for peck rates as a function of being engaged in cover (1) or not (0) were generated using a binomial general linear model.

|  | *Estimate* | *SE* | *Z* | *P* |
| --- | --- | --- | --- | --- |
| Intercept | -1.988 | 0.058 | -34.174 | < 0.001 |
| Engaged in cover (0:no, 1: yes) | 0.526 | 0.194 | -7.856 | < 0.001 |

## DISCUSSION

In this study, we provide the first evidence for cooperative breeding in the vulturine guineafowl (*Acryllium vulturinum*). Vulturine guineafowl groups contain multiple females (we observed up to 8 breeding females out of 13 adult female group members) that can attempt to reproduce within the same breeding season, meaning that they are also clearly plural breeders. This makes them similar, in terms of social and reproductive system, to the sympatric superb starling (*Lamprotornis superbus*) (Rubenstein, 2006). We found that carers provide the majority of the care and pay a foraging cost when doing so. As in most avian cooperatively breeding birds, we found that caring associations and caring interactions were very male biased. Overall, vulturine guineafowl have the common hallmarks of plural cooperatively breeding species.

The predominant hypothesis explaining the evolution of cooperative breeding behaviour is inclusive fitness (Hamilton, 1964). By helping, individuals can gain direct fitness benefits through routes including parentage of offspring (Richardson et al., 2002), territory inheritance (Kingma, 2017), group augmentation (Ligon & Ligon, 1978; Wright et al., 2010) or ‘pay-to-stay’ benefits (Wong & Balshine, 2011). In contrast to these direct routes, indirect fitness benefits may be accrued by helping kin, and can be through enhanced offspring number (Blackmore & Heinsohn, 2007) or survival (Hatchwell et al., 2004), as well as increased parental survival (Downing et al., 2021) or reproductive rate (Russell, 2003). As male birds are typically philopatric (Greenwood, 1980), non-breeding males (unlike the dispersing females) are likely to be related to breeders and their offspring, and may thereby gain indirect fitness by helping (Dickinson & Hatchwell, 2004). In precocial species where breeders invest considerably in reproduction, additional offspring care from such group members could reasonably generate indirect fitness benefits—such as greater offspring survival or enhanced parental current or future reproductive investment—that outweigh the costs of helping.

Cooperatively breeding vulturine guineafowl conform to the hypothesised patterns of cooperative breeding in birds. Carers are likely to increase the reproductive success of the female or enhance the female’s survival through reducing her need to care directly for the chicks. Specifically, we found that the female only provides a small proportion of all the chick caring interactions, which is likely to be important for her to regain the body condition she lost through laying and attending to the nest. Further, as female vulturine guineafowl may breed twice per year—as we found with several females being successful in both May 2020 and November 2020 (Table 1)—additional caring may be key to females regaining body condition in time for the next reproductive opportunity by reducing the inter-birth interval (Ridley & Raihani, 2008). Thus, our study provides further support for a number of general hypotheses surrounding cooperative breeding.

There is considerable variation in the structure of cooperatively breeding groups among species, ranging from a single breeding pair with associated helpers to multiple breeding units, which may be polygamous or polyandrous (Koenig & Dickinson, 2016; Stacey & Koenig, 1990). We still have no clear understanding of the drivers that give rise to multiple breeding units in the vulturine guineafowl, where multiple breeding pairs can form within a group during the breeding season and apparently non-breeding group members provide care to the offspring after hatching. This may be similar to golden lion tamarins (*Leontopithecus rosalia*), which have multiple breeding individuals in which adults care for offspring regardless of how many offspring there are in a cooperative polyandrous group (Dietz & Baker, 1993; Goldizen, 1989). However, in contrast to the golden lion tamarins, we found clear evidence for specialisation among non-breeding group members in term of which female they helped—likely their mothers. Plural breeding may be facilitated by breeding during a wet season, coinciding with a temporary increase in resource abundance (both food and safe nesting sites) that allow many individuals to reproduce at the same time. Our study also highlights the variance that can emerge in terms of who is a successful breeder, with many clutches depredated during incubation or early in the life of chicks. We ruled out dominance as a determinant for the breeding success in females, because the structure of the female dominance hierarchy is not very clear relative to that of males (Dehnen et al., 2022). However, more data are needed to more explicitly evaluate the link between breeding and dominance, and whether factors such as experience might drive variation in nesting rates and nest success among females.

In this study, we identified four juvenile-directed cooperative behaviours by non-parents, including babysitting (BBS), chick feeding (CFD), covering the chicks under the wings (COV), and within-group chick guarding behaviour against other adults (GRD). These are consistent with cooperative breeding behaviours exhibited in other precocial species (DuPlessis et al., 1995). For example, cooperative breeding behaviours of Kalij pheasants (*Lophura leucomelanos*) include caring for chicks, vigilance against predators and agonistic interactions among conspecific intruders (Zeng et al., 2016). Similarly, helpers in the buff-throated partridge (*Arborophila brunneopectus*) care for the chicks by identifying food and remaining vigilant against intruders (B. Wang et al., 2017). Such behaviours are clearly beneficial to chicks but may not be immediately obvious to observers and typically require following groups at small distances to make close observations. Doing so in our study was made substantially easier by the focal social group being habituated to our presence and due to their home range being centred within a fenced area that was safe to walk in.

The associates of each brood were predominantly male group members, and they typically provided more care than the mother did. These results are consistent with those reported in other cooperatively breeding Galliformes. One exception in our data was the second breeding season, although this season was notable for having many failed nests, which could have stimulated care from females. Why males attend to broods more than females in vulturine guineafowl, given that subadult females can remain in their natal territory for several years before dispersing (Klarevas-Irby et al., 2021), remains unknown, but this pattern has been observed widely in other species (Green et al., 2016). Further, we found that while some females were detected as significant associates of the brood, male carers provided nearly 98% of all the chick feeding interactions (excluding those from the mothers). This is consistent with the grey-crowned babbler (*Pomatostomus temporalis*), where only the number of male helpers increased reproductive success (Blackmore & Heinsohn, 2007).

By studying the social network of individual carers and broods, we showed that carers appeared to be generally brood specific. Among these, we found several lines of evidence supporting the hypothesis that vulturine guineafowl carers include non-breeding males. This includes subadult males not only associating significantly with their mother’s brood, but also providing care before they are fully grown (e.g., CT311, Figure 2). Others (e.g., CT317, a male that cared for YOBK’s brood in the second and third seasons) were part of the brood that their social mothers cared for before being observed providing care themselves. Further, the significant associates with each brood are relatively consistent over years. For example, YKOY was not detected as a significant carer in the first season, when GOOP did not successfully breed, but was significantly associated with her brood in the following two seasons. Thus, care is not given randomly within the group, and a substantial proportion of the care that we observed came from individual males that are unlikely to have any paternity with the brood.

A consistent pattern that emerged across all three seasons is that mothers did not provide the majority of care to their chicks. The large amount of help given by non-mother is perhaps unusual (Green et al., 2016). For example, in purple gallinules (*Porphyrio martinicus*), breeding adults provide the majority of the care to chicks, which are sub precocial (Hunter, 1987), while both male and female purple gallinules participate in incubation (Gross & Van Tyne, 1929). One reason why female vulturine guineafowl receive so much help in raising their offspring may be the high cost they pay during incubation—meaning that recovering their body condition might compromise the amount of care they can provide to the current brood, which is then offset by the care provided by other individuals. Further, care involves not only food provisioning, vigilance against predators and agonistic behaviours against intruders (Clutton-Brock & Manser, 2016) but also learning so as to enhance foraging skills (Cant et al., 2016; Heinsohn, 1991).

Cooperative breeding behaviours are likely to be costly to carers (Covas et al., 2022; Cram et al., 2015; Guindre-Parker & Rubenstein, 2018a, 2018b; Mendonça et al., 2020). For example, in meerkats (*Suricata suricatta*), helpers lose weight when they participate in cooperative breeding activities, such as feeding the young (Russell, 2003). Similarly, in white-winged choughs (*Corcorax melanorhamphos*), helpers lose weight when performing incubation, in addition to costs they incur by choosing to remain in their natal territory (Heinsohn & Cockburn, 1994). Our study adds to this body of evidence, demonstrating that individuals spend less time foraging when engaging in cooperative breeding behaviours relative to other group members that do not provide care. While we report that vulturine guineafowl carers incur some foraging costs, our data do not yet allow us to quantify the total extent of the costs they bear.

Further behavioural studies on cooperative breeding are needed in non-passerines, species with precocial young, and plural breeders. Specifically, we suggest that future research investigates what determines the relative reproductive success of males and females, especially in species where nest predation is high, which may result in hatching success to be relatively random among females. Such insights are important for understanding how indirect fitness might be gained via paternal routes, which are usually linked with much greater kinship uncertainty. Finally, further studies are needed on the reproductive behaviours of precocial species, where female investment (or lack of help received) during incubation, together with high levels of sociality, may be an indicator of which species breed cooperatively.

## ETHICS

Ethical approval was granted by the Max Planck Society’s Ethikrat Committee. Data were collected with permission from, and in collaboration with, the Kenya National Science and Technology Council (NACOSTI/P/16/3706/6465), the National Environment Management Authority (NEMA Access Permit NEMA/AGR/68/2017), the Kenya Wildlife Service, the National Museums of Kenya, Dr. Peter Njoroge, and the Mpala Research Centre.

## ACKNOWLEDGEMENTS

We thank the vulturine guineafowl team members, especially Wismer Cherono and Deborah Molitor for assistance with data collection and the Farine lab for discussions about the study.

## SUPPLEMENTARY MATERIALS

**Table S1:**
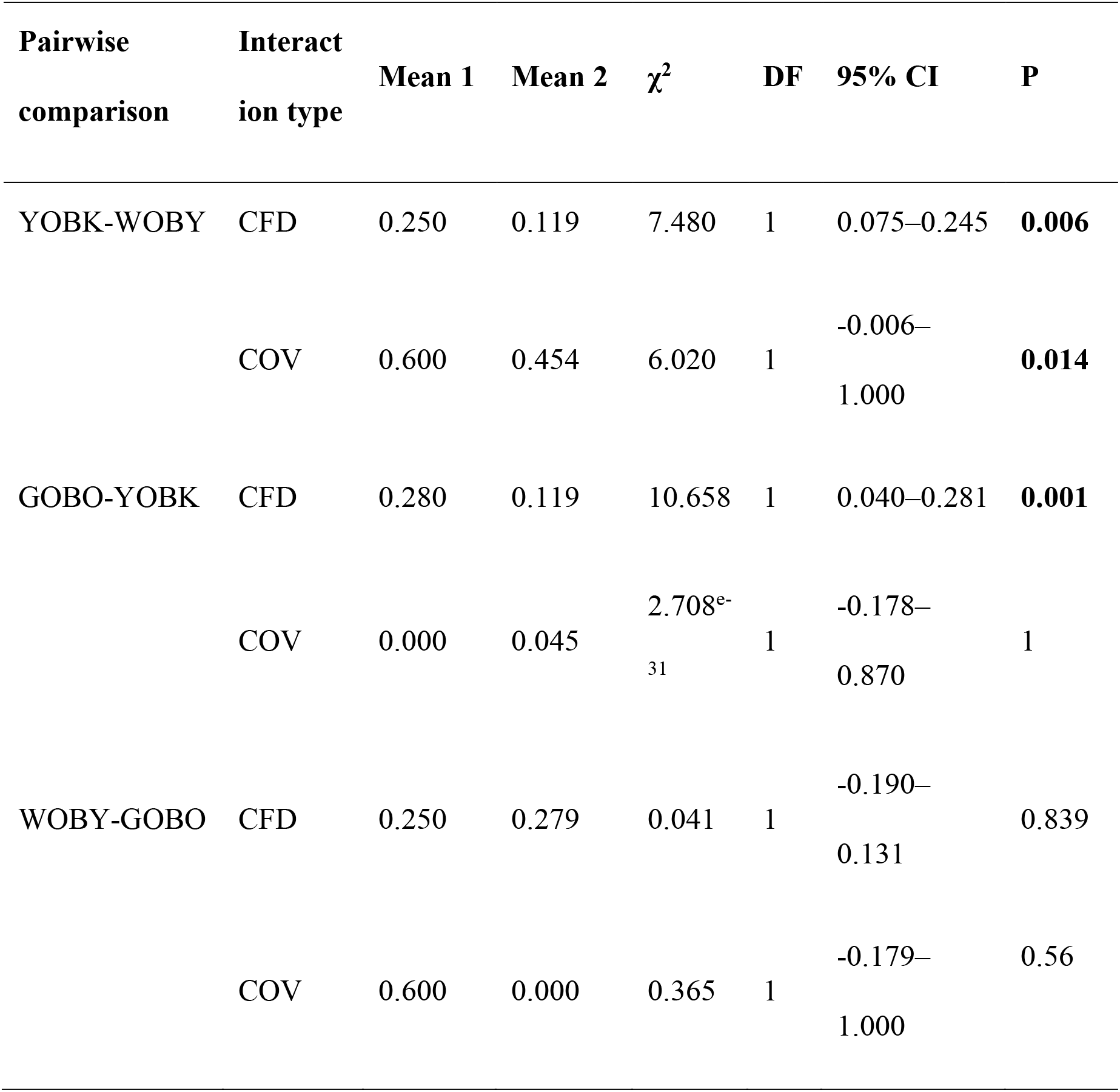
Overview of two-sample tests for equality of proportions, comparing the proportion of help given by the mother relative to the help given by helpers across the three broods in season 1. Significant values are shown in bold.

| Pairwise<br>comparison | Interact<br>ion type | Mean 1 | Mean 2 | $\chi^2$ | DF | 95% CI | P |
| --- | --- | --- | --- | --- | --- | --- | --- |
| YOBK-WOBY | CFD | 0.250 | 0.119 | 7.480 | 1 | 0.075–0.245 | **0.006** |
|  | COV | 0.600 | 0.454 | 6.020 | 1 | -0.006–<br>1.000 | **0.014** |
| GOBO-YOBK | CFD | 0.280 | 0.119 | 10.658 | 1 | 0.040–0.281 | **0.001** |
|  | COV | 0.000 | 0.045 | 2.708<sup>e-31</sup> | 1 | -0.178–<br>0.870 | 1 |
| WOBY-GOBO | CFD | 0.250 | 0.279 | 0.041 | 1 | -0.190–<br>0.131 | 0.839 |
|  | COV | 0.600 | 0.000 | 0.365 | 1 | -0.179–<br>1.000 | 0.56 |

**Table S2:** Overview of two-sample tests for equality of proportions, comparing the proportion of help given by the mother relative to the help given by helpers across the three broods in season 2. Significant values are shown in bold.

| Pairwise<br>comparison | Interact<br>ion type | Mean 1 | Mean 2 | $\chi^2$ | DF | 95% CI | P |
| --- | --- | --- | --- | --- | --- | --- | --- |
| YOBK-GOOP | CFD | 0.067 | 0.2333 | 4.1074 | 1 | -0.316-0.0178 | **0.042** |

**Table S3:** Overview of two-sample tests for equality of proportions, comparing the proportion of help given by the mother relative to the help given by helpers across the three broods in season 3. Significant values are shown in bold.

| Pairwise<br>comparison | Interac<br>tion<br>type | Mean 1 | Mean 2 | $\chi^2$ | DF | 95% CI | P |
| --- | --- | --- | --- | --- | --- | --- | --- |
| YOBK-RKGK | CFD | 0.339 | 0.068 | 61.489 | 1 | 0.205-0.337 | **4.450<sup>e-15</sup>** |
|  | COV | 0.333 | 0.000 | 2.341 | 1 | -0.0719-0.739 | 0.126 |
| YOBK-GOOP | CFD | 0.168 | 0.068 | 15.064 | 1 | 0.0544-0.146 | **0.001** |
| YOBK-GAGA | CFD | 0.100 | 0.068 | 1.923 | 1 | -0.010-0.073 | 0.166 |

**Figure S1.**
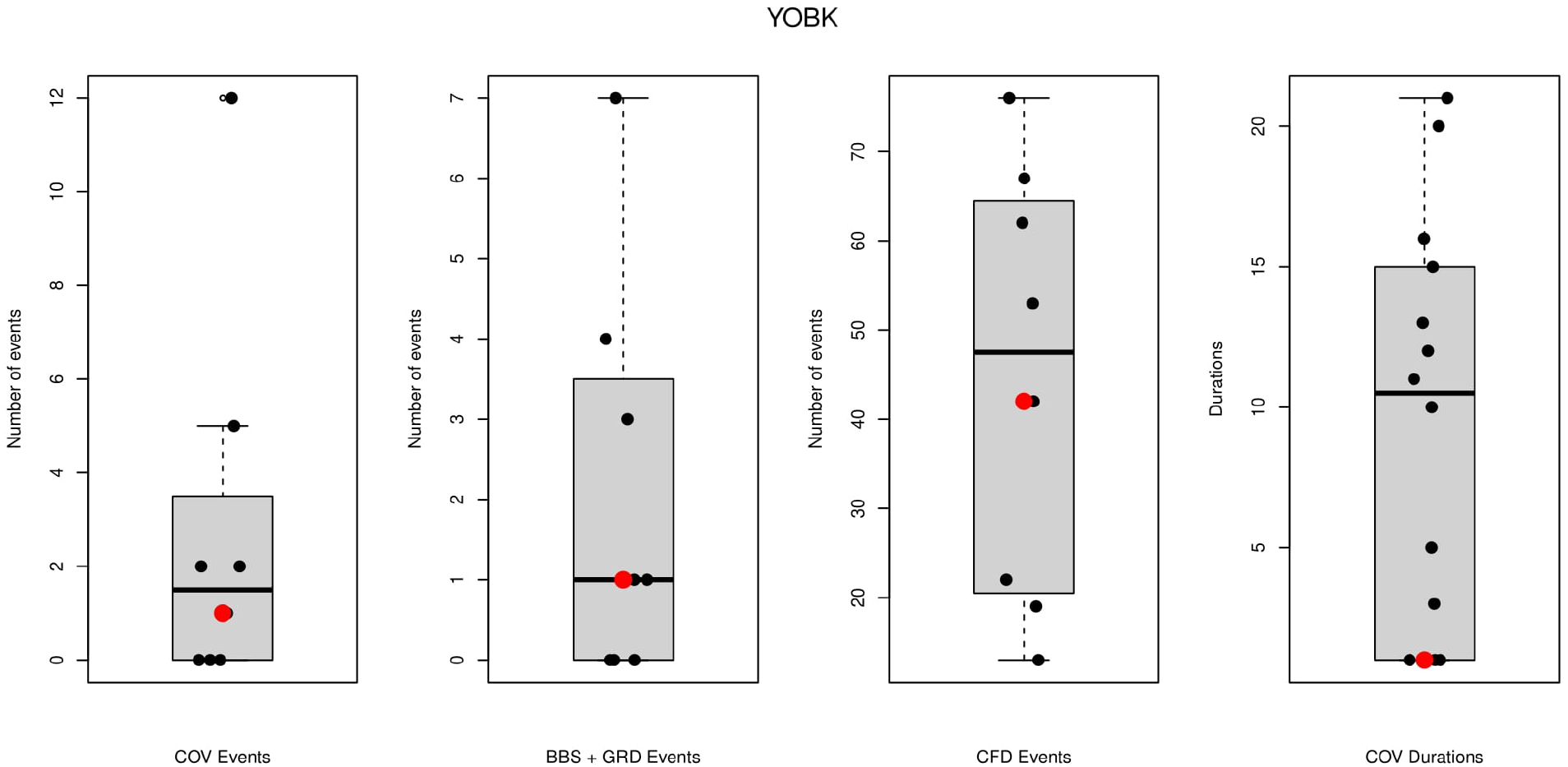
Amount of help given by the mother (red points) and helpers (black points) for the YOBK brood in season 1.

**Figure S2.**
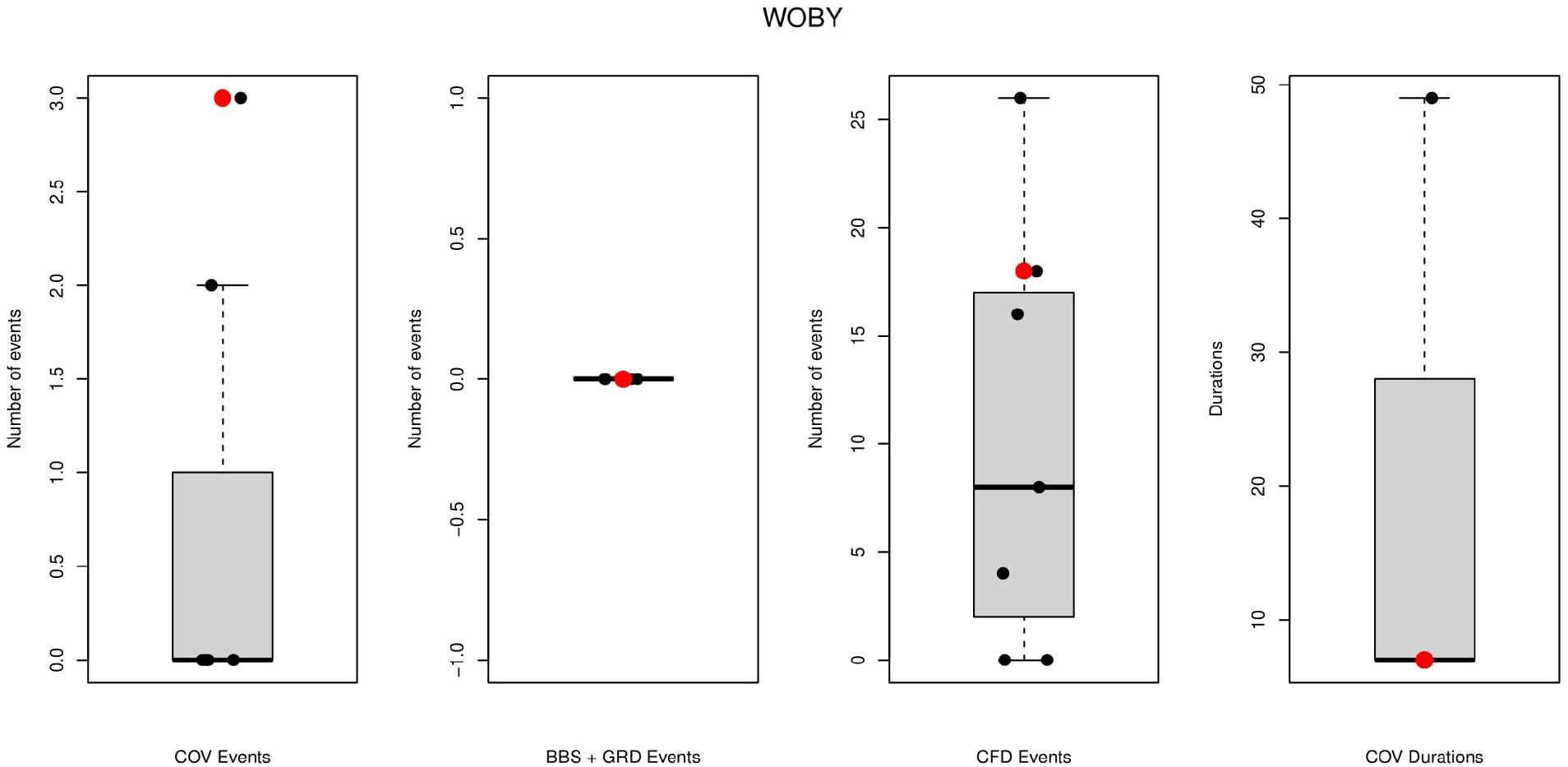
Amount of help given by the mother (red points) and helpers (black points) for the WOBY brood in season 1.

**Figure S3.**
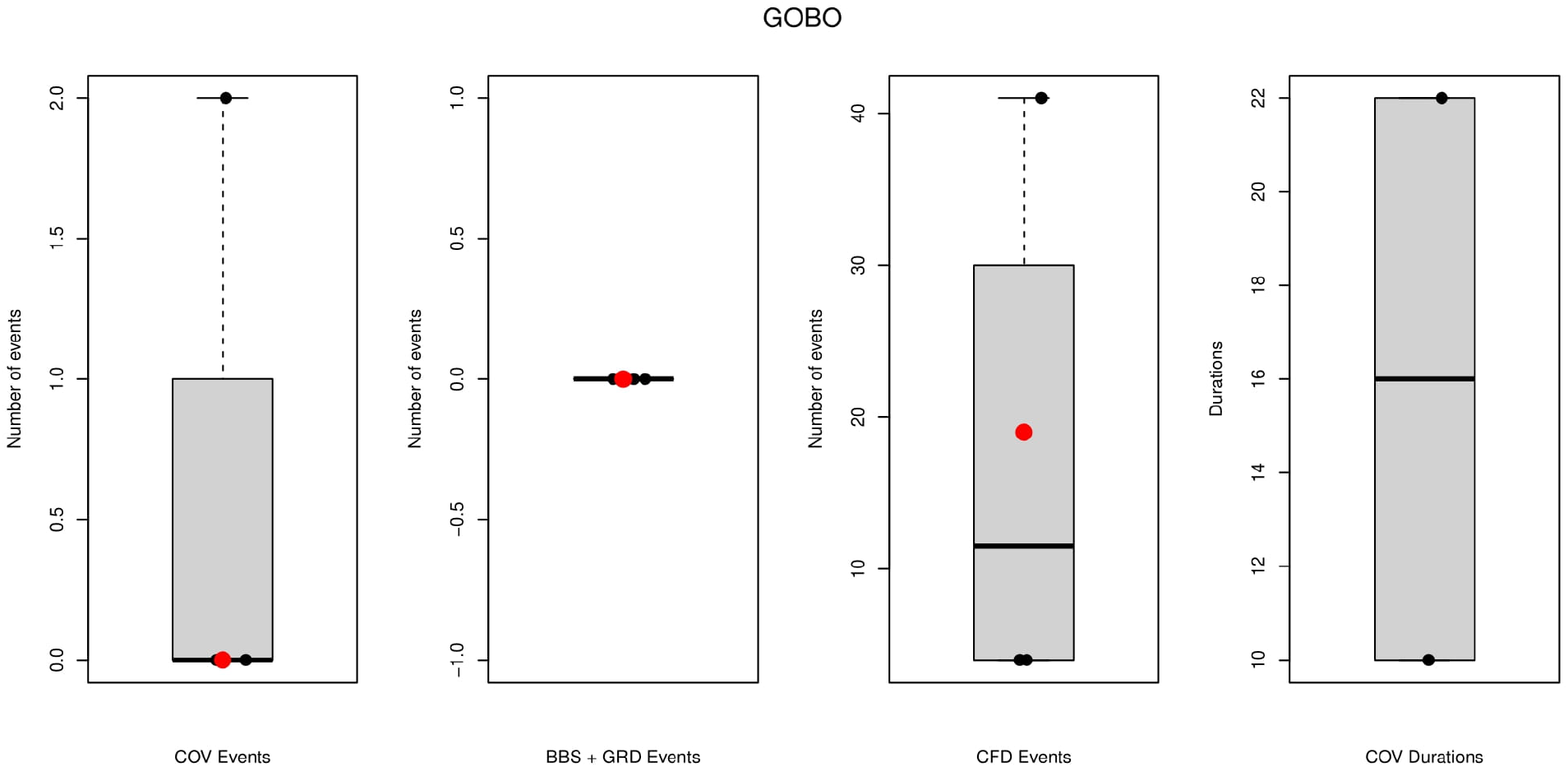
Amount of help given by the mother (red points) and helpers (black points) for the GOBO brood in season 1.

**Figure S4.**
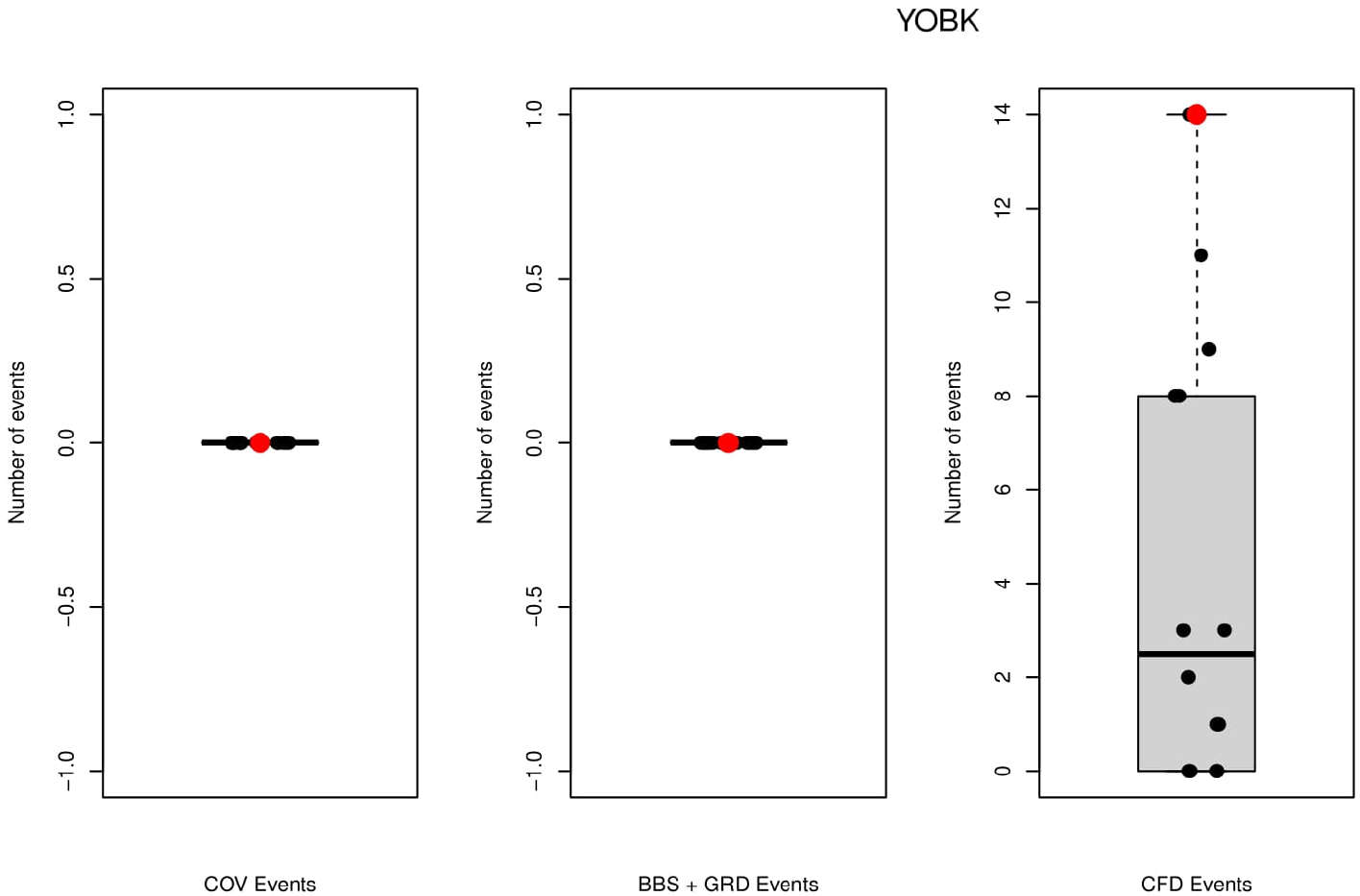
Amount of help given by the mother (red points) and helpers (black points) for the YOBK brood in season 2.

**Figure S5.**
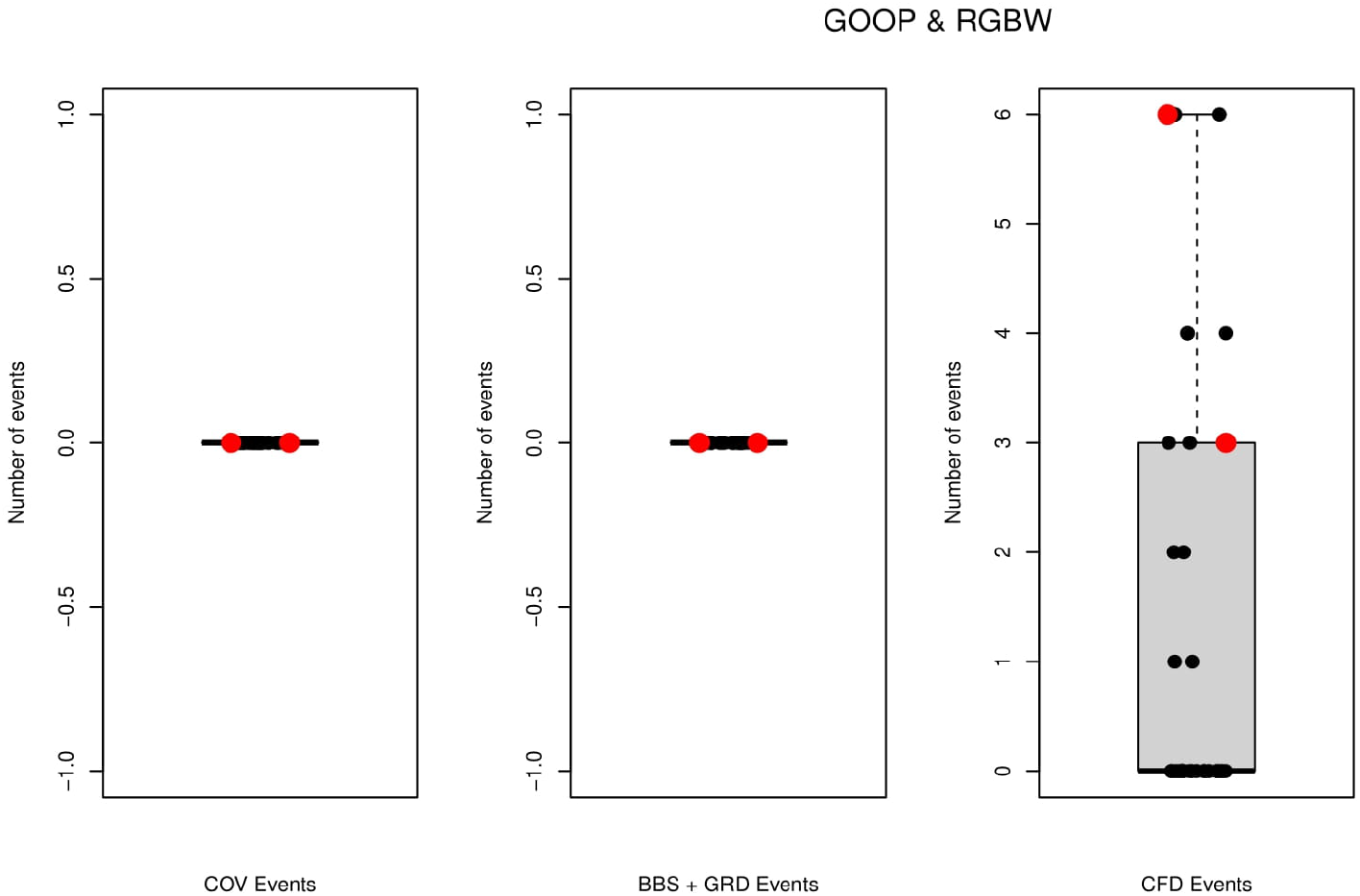
Amount of help given by the mother (red points) and helpers (black points) for the GOOP (and chicks from RGBW) brood in season 2.

**Figure S6.**
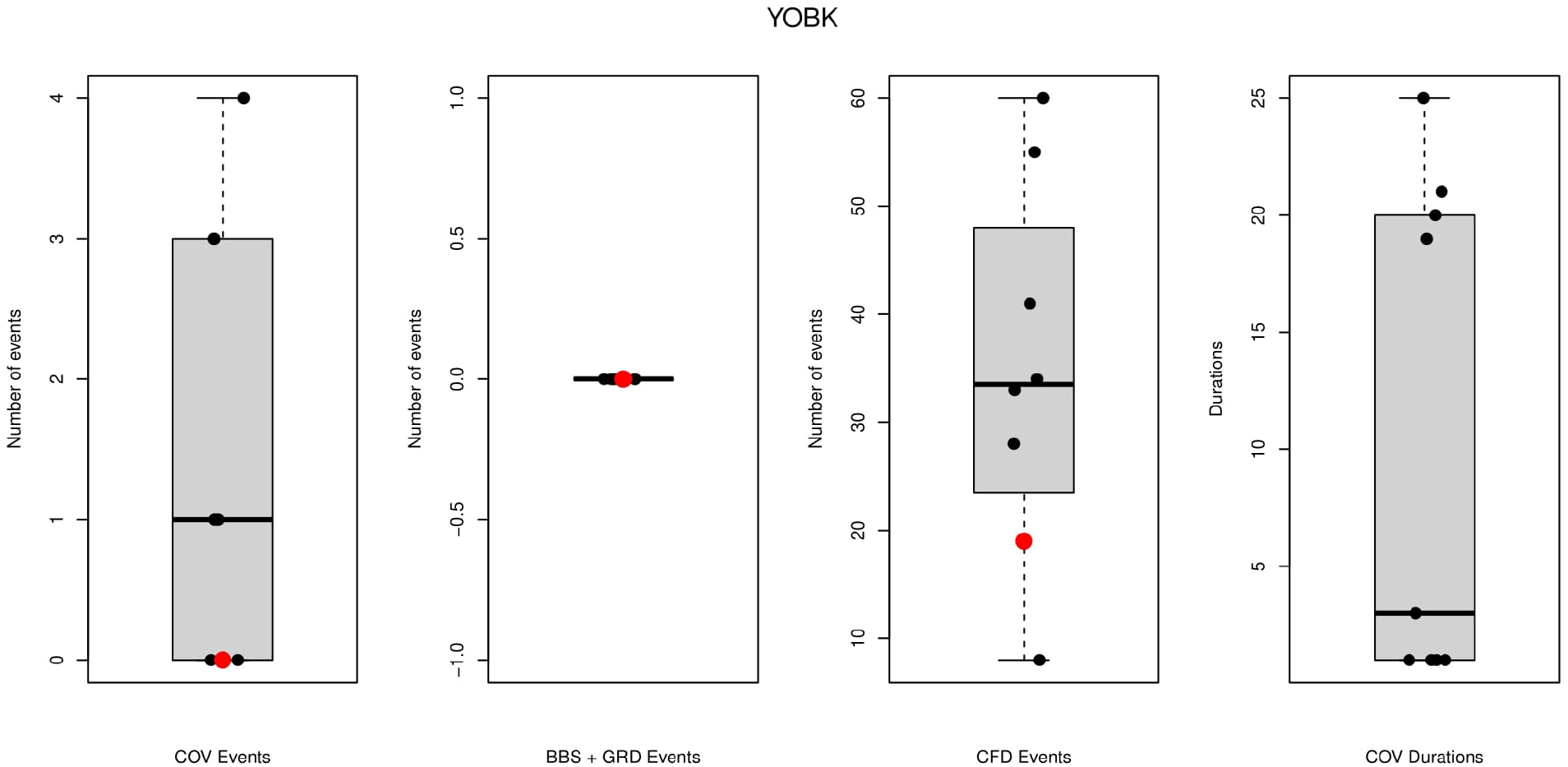
Amount of help given by the mother (red points) and helpers (black points) for the YOBK brood in season 3.

**Figure S7.**
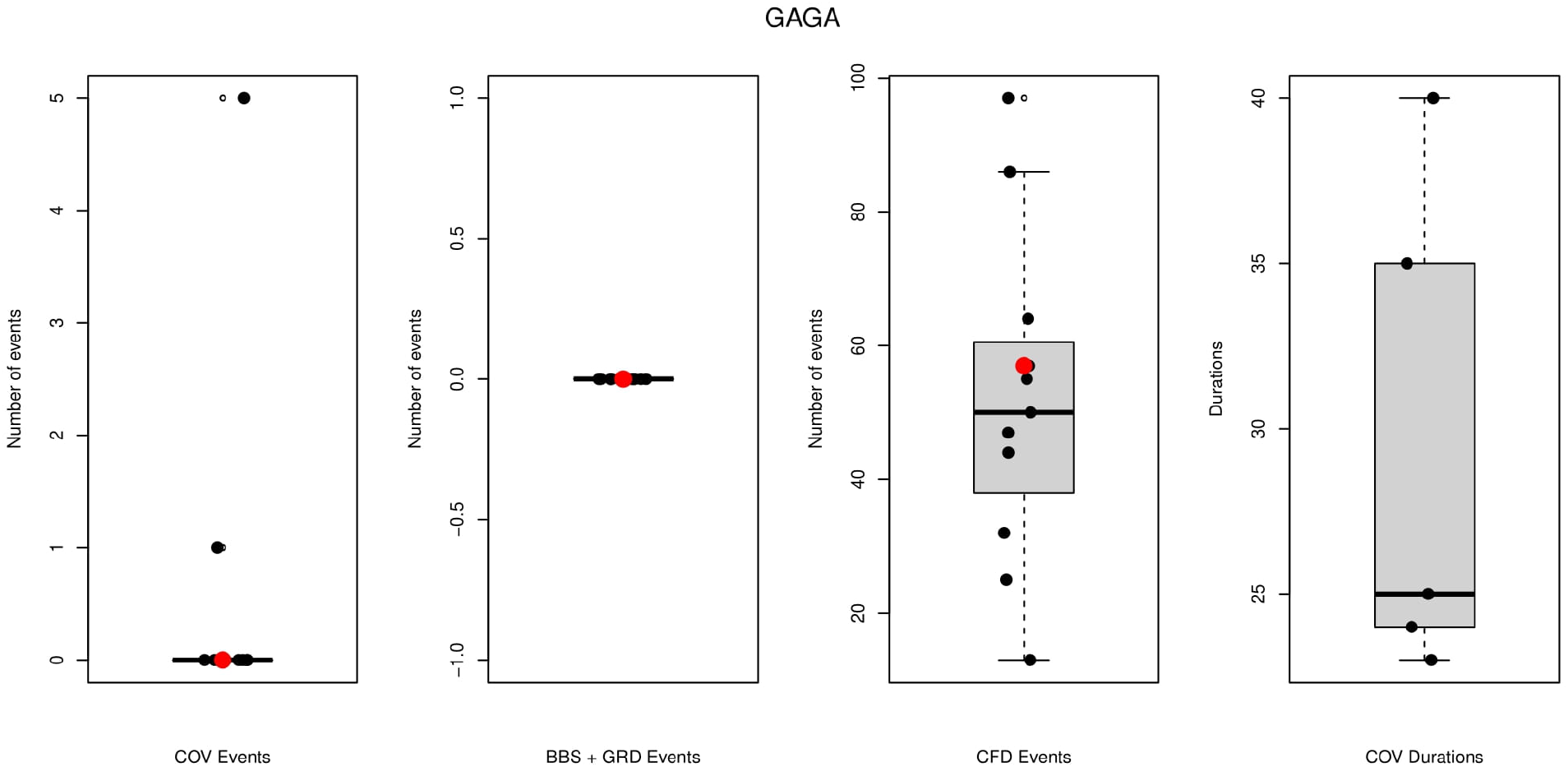
Amount of help given by the mother (red points) and helpers (black points) for the GAGA brood in season 3.

**Figure S8.**
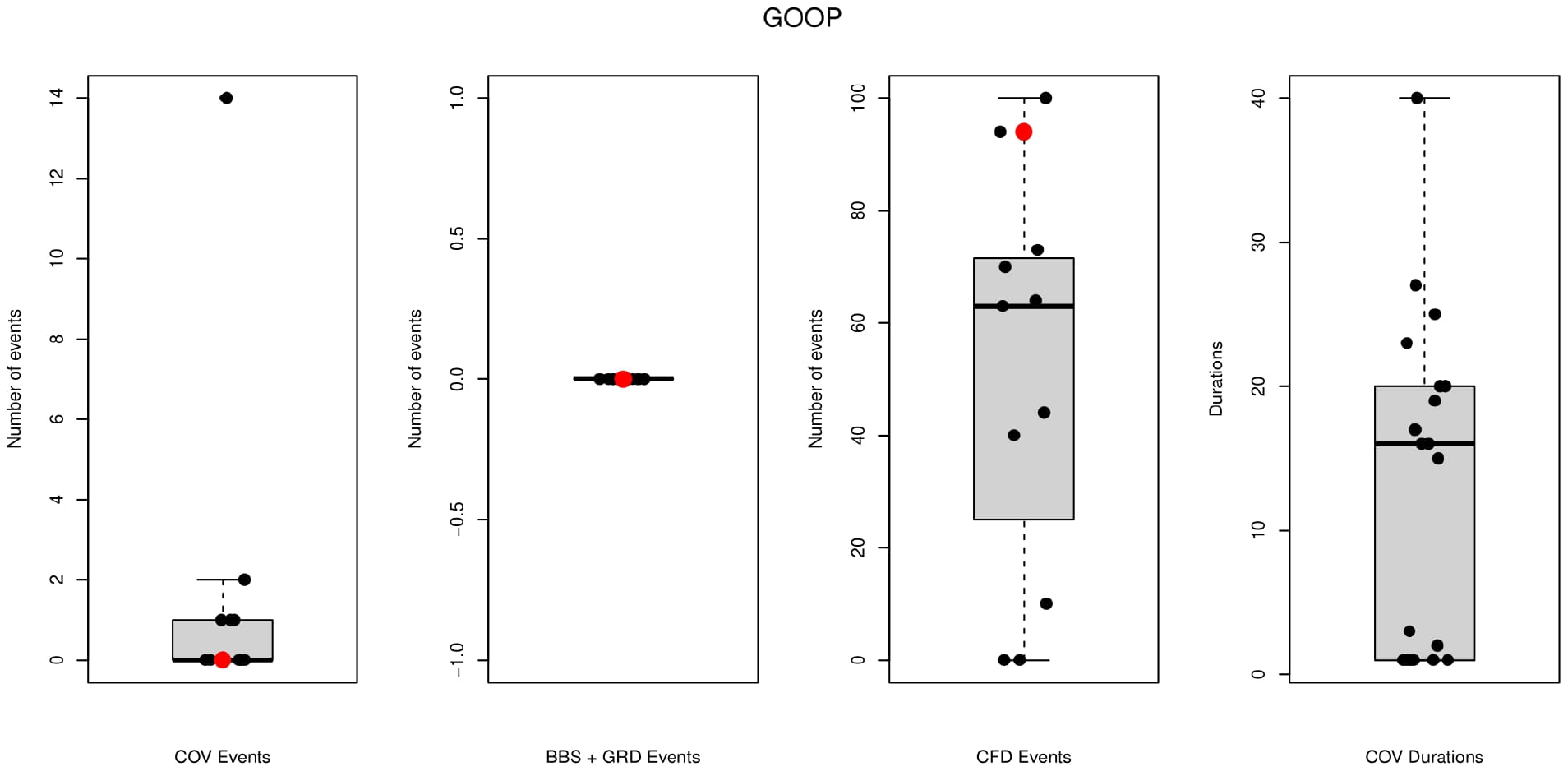
Amount of help given by the mother (red points) and helpers (black points) for the GOOP brood in season 3.

**Figure S9.**
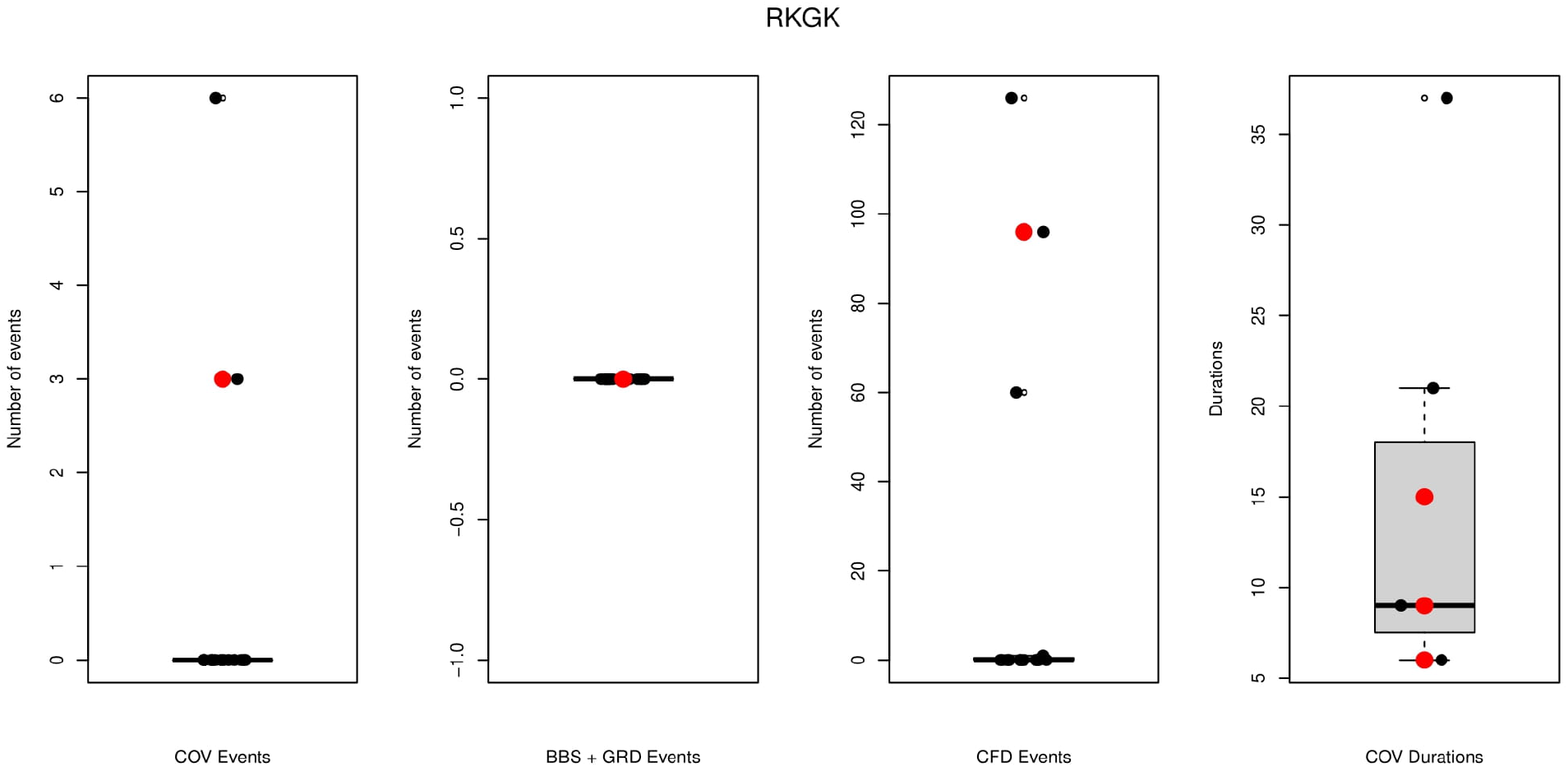
Amount of help given by the mother and helpers for the RKGK brood in season 3.

